# Addressing the “open world”: detecting and segmenting pollen on palynological slides with deep learning

**DOI:** 10.1101/2025.01.05.631390

**Authors:** Jennifer T. Feng, Sandeep Puthanveetil Satheesan, Shu Kong, Timme H. Donders, Surangi W. Punyasena

## Abstract

In the “open world”, categorical classes are imbalanced, test classes are not known *a priori*, and test data are captured across different domains. Paleontological data can be described as open-world, as specimens may include new, unknown taxa, and the data collected, such as measurements or images, may not be standardized across different studies. Fossil pollen analysis is one example of an open-world problem in paleontology. Pollen samples capture large numbers of specimens, including not only common types but also rare and even novel taxa. Pollen is diverse morphologically and features can be altered during fossilization. Additionally, there is little standardization in the methods used to capture and catalog pollen images and most collections are mounted on microscope slides. Therefore, generalized workflows for automated pollen analysis require techniques that are robust to these differences and can work with microscope images. We focus on a critical first step, the detection of pollen specimens on a palynological slide and review how existing methods can be employed to build robust and generalizable analysis pipelines. First, we demonstrate how a **mixture-of-experts approach** – the fusion of a general pollen detector with an expert model trained on minority classes – can be used to address taxonomic biases in detections, particularly the missed detections of rarer pollen types. Second, we demonstrate the efficiency of **domain fine-tuning** in addressing domain gaps – differences in image magnification and resolution across microscopes, and of taxa across different sample sources. Third, we demonstrate the importance of **continual learning workflows,** which integrate expert feedback, in training detection models from incomplete data. Finally, we demonstrate how **cutting-edge segmentation models** can be used to refine and clean detections for downstream deep learning classification models.

**Non-technical Abstract:** Fossil pollen analysis presents challenges that result from the complexity of the “open world”. It is not possible to anticipate all the taxa that will be encountered in geologic samples. Each sample has the potential to produce new species and introduce specimens with unique shapes and sizes. Pollen embodies diverse morphologies, its appearance can be altered depending on the conditions under which it was fossilized, and there is little standardization in the microscopy equipment used to image and catalog pollen images. AI applications in palynology need to address this complexity. We applied four machine learning methods (mixture-of-expert models, domain fine-tuning, continual learning, and foundation models for segmentation) to fossil pollen data to demonstrate their effectiveness in the detection and isolation of pollen specimens on a pollen sample slide.

## Introduction

The “open world” describes uncontrolled operational environments in machine learning (Liu et al. 2019; Joseph et al. 2021; Bendale and Boult 2015). It stands in contrast to classic supervised learning, where all the test classes are known *a priori* and have been introduced in training. Fossil pollen analysis represents a paleontological example of an open-world problem. Automated detection and classification of specimens has been a long sought-after goal given the high level expertise and time required for traditional analysis (Langford et al 1990). Specimens often represent new, undiscovered species. Abundances naturally follow a long-tailed distribution and as a result many taxa are rare and training examples are unbalanced. Images of the same taxon can vary widely, depending on the microscope, magnification, and imaging or preparation techniques. Specimens can have variable levels of preservation from sample to sample and locality to locality. Developing machine learning models that can generalize across the range of variability in pollen diversity, abundance, preservation, preparation, and imaging requires adopting workflows that can succeed in open-world environments.

Fossil pollen isolated from geologic sediments for paleoecological or biostratigraphic analysis are typically mounted on microscope slides (Traverse 2007). The widespread availability of slide scanning microscopes means that entire slides can be imaged quickly and efficiently (Punyasena et al. 2022). These scans are capable of capturing the entirety of a pollen slide - both the area of a coverslip and multiple focal planes. This produces a fully three-dimensional representation of the pollen sample (Punyasena et al. 2022).

Accurate, consistent detection of pollen among other organic debris in these slide scans is a critical step in automating visual pollen identifications. Detection in this context refers to the automated location of a pollen specimen within the X, Y, and Z coordinates of a slide scan. Detection is needed at two discrete stages of automated workflows. Detected pollen grains can be labeled by experts to efficiently produce training and validation data for the development of pollen classification models. Detection is also needed when applying these classification models to new samples.

Developing fully automated detection pipelines requires adapting models to the uncertainties inherent to open-world problems. We demonstrate four machine-learning solutions for addressing common limitations in automated pollen detection. Taken together, these approaches provide effective strategies for constructing robust, generalized detection models that can be applied to a wide range of palynomorphs and other paleobiological data.

We first demonstrate how the **mixture-of-experts technique** can address taxonomic bias in pollen detections. The morphological diversity and long tail of rare species encountered in pollen samples, particularly those from the tropics, lead to detectors that are potentially biased toward the most common and distinctive morphological types. False negatives (pollen that is missed by the detector) pose a greater problem than false positives (non-pollen objects identified as pollen by the detector), as false positives can be removed at a later stage of the analysis, while bias toward false negatives will affect downstream estimates of proportional abundance. The solution is to train an expert model on small, difficult-to-detect taxa and fuse it with a more general pollen detector. This technique is used frequently in top-ranked detection methods in public machine-learning challenges (Huang et al. 2020; Guo et al. 2019; Akiba et al. 2018).

We next demonstrate how data efficient **fine-tuning** can be in transfer learning across new domains. Differences among pollen samples or images result from differences in the microscopes and objectives used, localities or ages represented, or preparation techniques applied. Detection models trained in one domain will have reduced accuracy when applied to a new domain due to differences in color, brightness, and resolution, or the taxonomic or taphonomic differences between samples. Fine-tuning previously trained models with training data from a new domain can quickly produce more generalized models. The approach is widely used in machine learning (for reviews, see Zhuang et al. 2020; Pan and Yang 2010).

Thirdly, we show how workflows for **continual learning** using human-in-the-loop annotation can address the problem of incomplete training data. In fossil pollen analysis, as in many other areas of paleobiological research, there is a high probability of encountering new taxa with each new sample. We inevitably begin with insufficient training data because it is not possible to curate images of all possible types. Incorporating expert feedback through human-in-the-loop annotation leverages trained models to annotate new, unlabeled data. Experts verify or revise low confidence detections and these new annotations are used to further fine-tune detection or classification models (Zhou et al. 2017; Adhikari and Huttunen 2021; Wang et al. 2020; Wu et al. 2022; Kirillov et al. 2023). The approach provides an efficient mechanism for improving pollen detection (as well as pollen classification) models over time.

Finally, we demonstrate how newly available **foundation segmentation models** (Meta’s Segment Anything Model 2, SAM-2) (Kirillov et al 2023; Ravi et al. 2024) can be incorporated into object detection pipelines of microscope images. Segmentation builds on detection results, producing masked images that follow an object’s outline. Cleanly segmented images improve future classification analyses by removing extraneous and potentially biasing information from an image background. SAM-2 takes a user prompt -- a point within an object, a defined region of interest, a mask, or a text description -- and returns a valid segmentation mask by combining an image encoder with a prompt encoder. The image embedding and encoded prompt are fed to a mask decoder, which predicts segmentation masks. SAM-2, which generalizes promptable image segmentation to video domains, is trained on a segmentation dataset of over one billion masks, one million still images, and 51,000 videos, allowing the model to provide unsupervised segmentation results that can surpass trained segmentation models (Ravi et al. 2024).

We review these four foundational and emerging techniques and demonstrate their application to palynological analysis in hope of encouraging and guiding other researchers in developing robust deep learning workflows. All Python code used to process slide scans into image stacks, annotate image stacks, train and evaluate detection models, and apply segmentation is available through our GitHub repositories and we provide Command Line Interface (CLI) Python software so that others can easily integrate these techniques into their own research (see Data Availability below).

## Methods

### Pollen Samples Source

We imaged pollen from two high-resolution sediment cores extracted from Laguna Pallcacocha, in El Cajas National Park, Ecuadorian Andes by Moy et al. (2002) and Hagemans et al. (2021). The first core (PAL 1999) extends through the Holocene (11,600 cal. yr. BP - present). The second parallel core (PAL IV) spans the 20th century (Hagemans et al. 2023). The vegetation surrounding the lake is dominated by cushion plants and patches of páramo shrubs (Hagemans et al. 2019). Volumetric samples were spiked with *Lycopodium clavatum* and prepared following standing protocols, including acetolysis and heavy liquid flotation (Faegri, Kaland, and Krzywinski 1989). Samples from PAL IV were mounted using glycerine and samples from PAL 1999 were mounted in a permanent mount.

### Imaging

We imaged 19 slides from the PAL IV core and three slides from the PAL 1999 core using two different microscopes (Fig. 1). PAL IV slides were imaged at 630x magnification (0.146 μm per pixel resolution) with a Leica DM 6000 B, an upright transmitted light microscope fitted with an automated XYZ stage and LAS 4.12 PowerMosaic software for the creation of image tile grids (Fig. 1B). One to three scans measuring 3698 μm x 2790 μm were taken from each slide. Each scan was composed of 400 image stacks of 1040 x 1392-pixel tiles, with seven or nine focal planes imaged at increments of 3 or 4 μm. A total of 2986 image stacks contained palynomorphs; these were used for model training. The three samples from the PAL 1999 core were imaged at 400x magnification (0.225 μm per pixel resolution) with a Hamamatsu NanoZoomer 2.0 HT slide scanning microscope. Nine focal planes were imaged in 3-μm increments for a focal depth range of 24 μm. Slide scans were processed into 1040 x 1392-pixel image stacks using the Bio-Formats Python library (Linkert et al. 2010). Three hundred randomly selected image stacks, approximately 5% of the total slide area, were exported from each slide.

**Figure 1.**
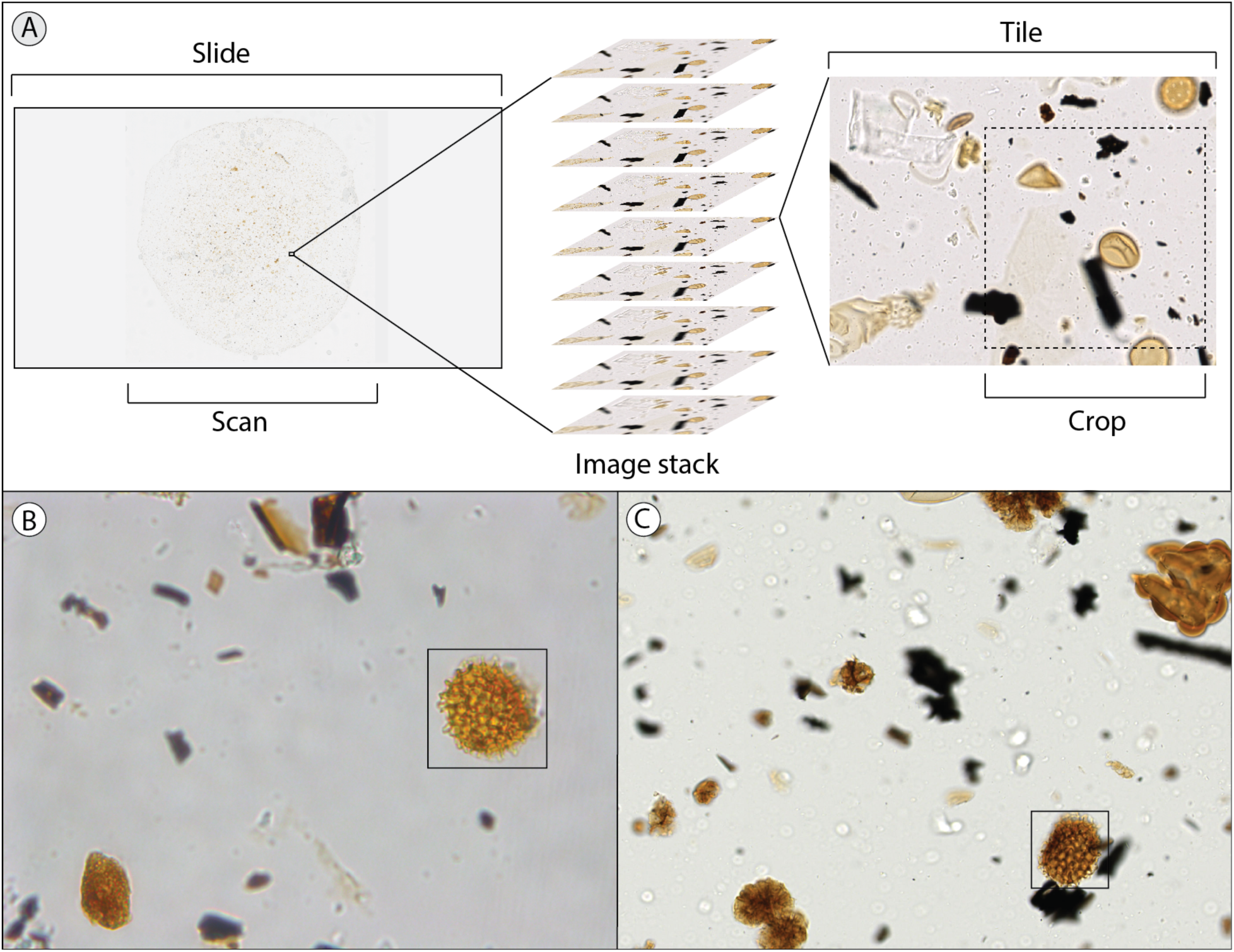
(A) Example of a slide scan (∼20.5 x 20.5mm), image stack (7 or 9 focal planes with 3 or 4 μm step size), image tile (1040 x 1392 pixels), and crop (800 x 800 pixels). Comparison of 1040 x 1392-pixel image tiles taken with a (B) standard upright microscope (0.146 μm/pixel) and (C) slide scanning microscope (0.225 μm/pixel). 40 x 40 μm boxes highlight *Lycopodium* spores in each image for comparison. Noticeable differences include color, brightness, and scale. The upright microscope domain was used in training the general pollen detection model and small-grain detection model and the slide scanning microscope domain was used for domain fine-tuning and continual learning.

### Annotation

For every annotated pollen grain or spore, we defined its XYZ coordinates using bounding boxes converted to inscribed circles (PAL IV images) or directly as circles (PAL 1999 images) on the plane of the image stack that captured the equatorial cross-section of the pollen grain or spore. When these annotations fell at the edge of the image, annotations were corrected manually to identify the center of the grain. We annotated the PAL IV images using a MATLAB script, modified from Punyasena et al. (2022), and annotated the PAL 1999 images using Labelme (Wada et al. 2021) and the Hamamatsu NanoZoomer NDP.view2 software. All three methods produced the same annotation metadata: a center and radius that defined the location of a pollen grain. We annotated 3191 PAL IV and 794 PAL 1999 specimens as one of 129 taxonomic types (Feng et al. 2025). The 18 most common palynomorphs accounted for 92% of the total dataset (Fig. 2). We excluded algae and fungal spores, but included plant spores from ferns and lycopods, such as *Huperzia* (Fig. 2P), *Isoetes* (Fig. 2Q), and the exote marker *Lycopodium clavatum* (Fig. 2R).

**Figure 2:**
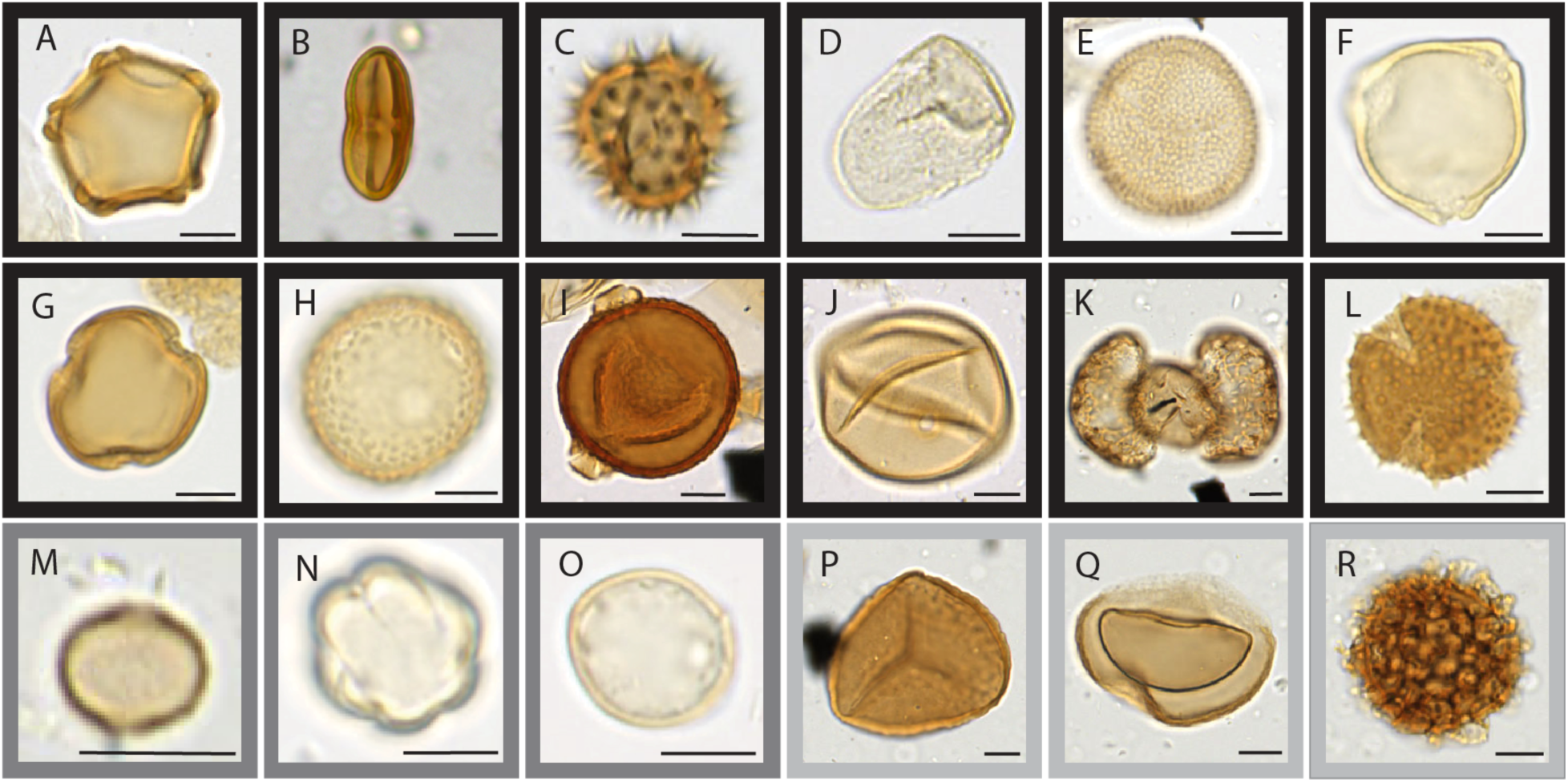
The 18 most common palynomorphs (>20 training examples, representing 92% of the total dataset): A) *Alnus*, B) Apiaceae, C) Asteraceae Tubuliflorae, D) Cyperaceae, E) *Hedyosmum*, F) *Myrica*, G) *Myrsine*, H) *Plantago*, I) *Polylepis* spp., J) Poaceae, K) *Podocarpus*, L) *Valeriana*, M) *Cecropia*, N) Melastomataceae, O) Urticaceae-Moraceae, P) *Huperzia*, Q) *Isoetes*, and R) *Lycopodium clavatum* (exote marker). Black borders indicate taxa with medium to large pollen grains (A – L), medium-gray borders indicate taxa with small grains (M – O), and light gray borders indicate plant spores included in the annotated dataset (P – R). Scale bars represent 10 μm.

### General Pollen Detection Model (GPDM) Architecture and Training

Convolutional neural networks (CNNs) have learnable parameters that are optimized to output the desired prediction for a given input. In our case, our input is image stacks from slide scans and our desired prediction is pollen detections. We used ResNet34 (He et al. 2016) as the backbone of our detection model and built a decoder on top of this backbone (Punyasena et al. 2022; Kong 2022). The decoder configured the output from the model as a detection map, with each pixel approximating a confidence score for predicted pollen detection.

There are three main steps in training a CNN. First, training data are forward-passed through the model to calculate the loss value, a measure of the model’s performance. Loss is the difference between a model’s outputs and ground-truth (i.e., taxa names and pollen grain bounding box coordinates), and its value is minimized through multiple training iterations.

Second, learnable parameters are updated according to an optimization algorithm (for a review of hyperparameter tuning, see Godbole et al. 2023). Finally, the model which produces the smallest loss on an independent validation set is selected. Once the model has been trained and the best model selected through validation (Fig. 3A – Step 1), the learnable parameters are fixed and the model is deployed to help analyze experimental data, e.g., using validation data to evaluate the model quantitatively (Fig. 3A – Step 2).

**Figure 3.**
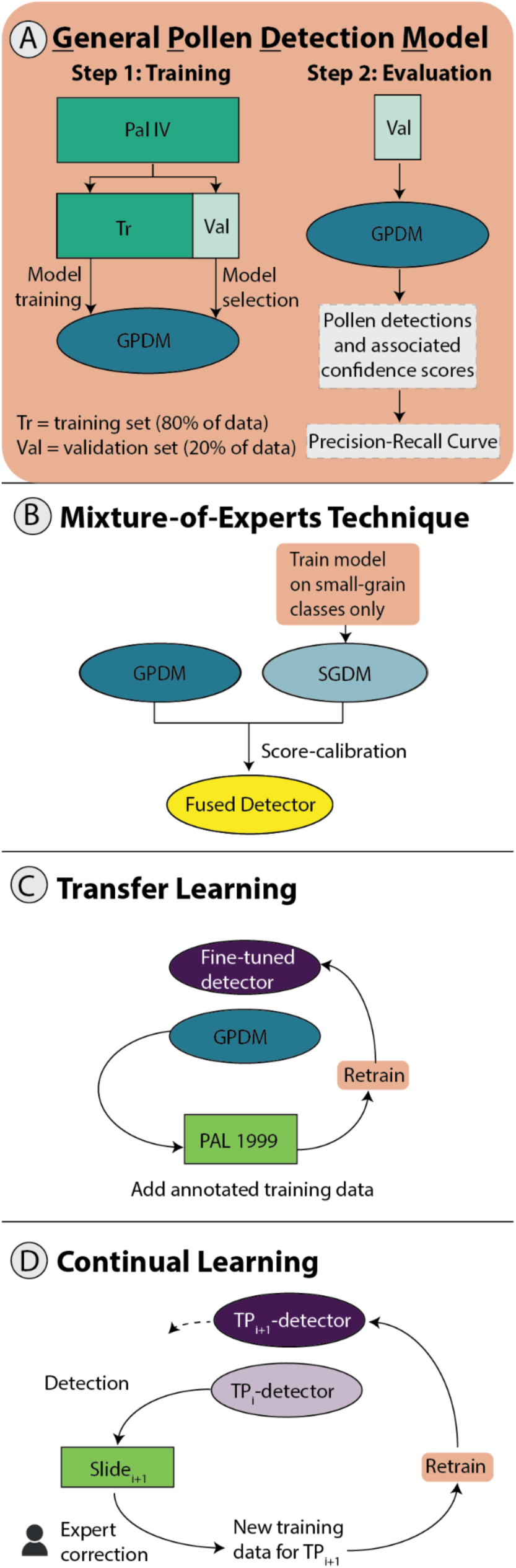
Setup for (A) the general pollen detection model (GPDM), (B) the mixture-of-experts technique, (C) transfer learning across imaging domains, and (D) continual learning with human-in-the-loop annotation. (A) shows the process of training the GPDM by splitting the annotated dataset into training and validation sets, used respectively for training the model and for model selection. The validation set was then passed through the model to obtain a list of detections and associated confidence scores. Using the detections, we drew a precision-recall curve for model evaluation. (B – D) are variations on (A). In (B), we trained a small-grain detection model (SGDM) on a subset of the PAL IV data, selecting only image stacks that contained Urticaceae-Moraceae, Melastomataceae, or Cecropia grains and revising the masks to only represent these taxa. The SGDM was then fused with the GPDM into a single pipeline. In (C), we fine-tuned the GPDM on slides from a new domain, PAL 1999. In (D), we implemented a continual learning workflow. We used Slide 1 to fine-tune the GPDM in TP_0_, producing the TP_1_-detector, then used the TP_1_-detector to detect pollen in image stacks from Slide 2. In the fine-tuning stage, experts manually verified detections so that the detections served as new training data to finetune detectors. The process was repeated continually in subsequent time steps.

We used the PAL IV dataset in developing our general pollen detection model (GPDM). We divided the annotated image stacks into a training set (80%) and a validation set (20%). We augmented the training data using random rotations and flips of the original image stacks. We used an image batch size of 4, trained the model for 30 epochs (where one epoch is the complete pass of the training dataset through the CNN), and set the base learning rate (the degree of model weight adjustment for each weight update) to 0.0005. The model was trained on 800 x 800-pixel crops of the original 1040 x 1392-pixel image tiles (Fig. 1A). During training, crops were taken randomly, and the location of the crop varied with each epoch. During evaluation, four overlapping crops were taken of each image stack, covering the entire image tile.

We created circular binary masks to indicate where pollen was present using our circular annotations. From the masks, we created distance transform masks where values for pixels inside each circle or partial circle represented the number of pixels to the nearest edge (Fig. 4B). Masks were used to train the model to recognize pixels in an image as “pollen” or “not pollen,” while the distance transform masks were used to train the model to recognize the centers of pollen grains. The loss score for each epoch of training was the sum of the detection loss and the distance transform loss.

**Figure 4.**
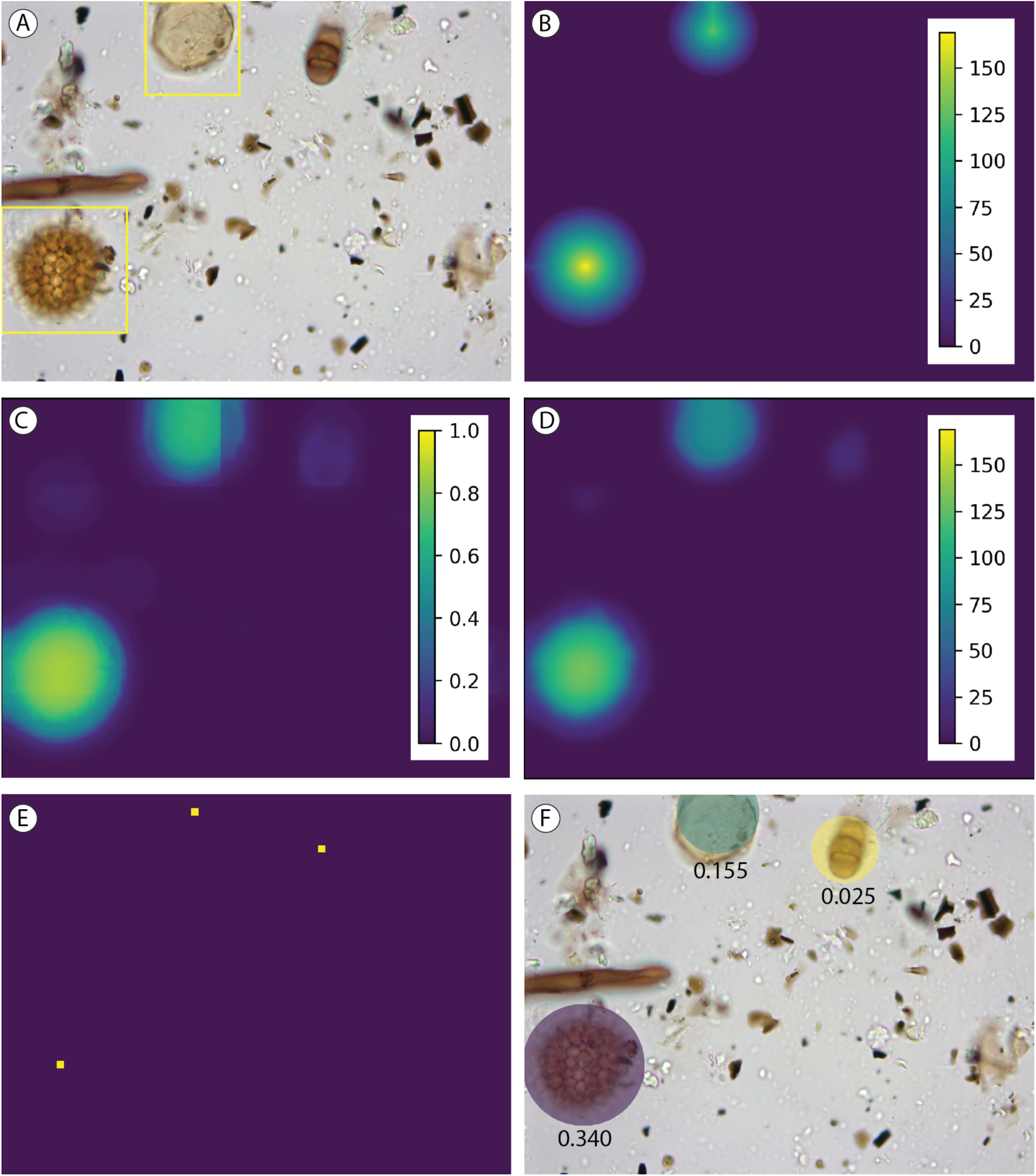
The detection workflow. (A) One plane of the image stack, overlaid with the original square annotations. (B) The ground-truth distance transform mask, created from the annotations. (C) The softmax layer, one of the model outputs. (D) The predicted distance transform mask, a second model output. (E) The predicted pollen grain centers, determined by calculating the peaks in the distance transform mask. (F) The detection mask, created using the predicted pollen grain centers and radii, overlaid on the image with confidence scores below each detection. The detection mask was thresholded at a confidence score of 0.025. Note that a single image is used solely to illustrate our workflow. In our study, training and evaluation images were not duplicated.

### Detection Model Outputs

For each image in an image stack, the detection model outputs softmax layers (normally distributed probabilities of whether a pixel is within a pollen grain) (Fig. 4C), and predicted distance transform layers of pollen pixels from the edge of a pollen grain (Fig. 4D). Local peaks in the distance transform layer identified the mass centers of pollen detections (Fig. 4E). We calculated the predicted radius by finding the largest connected component and fitting an ellipse. The predicted center and radius defined the binary detection mask (Fig. 4F). Softmax was used to determine the confidence of the detection (Punyasena et al. 2022).

### Detection Model Evaluation

We used confidence scores to rank and select model detections. Lowering the confidence threshold to increase detections introduced a tradeoff between precision and recall. Precision is the percentage of detections that are true positives. Recall is the percentage of the annotated pollen that is detected. The two metrics can be summarized as:

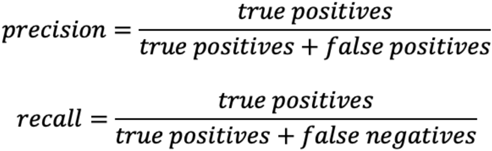

The area under the precision-recall curve captures the mean average precision (mAP), which we used to evaluate model performance (Everingham et al. 2010; Lin et al. 2014).

We used intersection over union (IoU, the area of overlap between two targets divided by the area of their union) (Padilla et al. 2021) to identify true positives, false positives, and false negatives. We established the IoU threshold (0.3) through visual assessment. If the IoU of a detection (Fig. 4F) and the nearest ground-truth annotation (Fig. 4A) was ≥ 0.3, the detection was considered a true positive, and if < 0.3, a false positive. If an annotation did not overlap with any detections with an IoU ≥ 0.3, it was a false negative. Within an image stack, there are multiple detections for the same grain. We used non-maximum suppression (NMS) to remove duplicates when the IoU of two detections was ≥ 0.3, retaining only the detection with the highest confidence score.

### Mixture-of-Experts

In our samples, small grains were only 6.8 % of the total dataset, so taxonomic bias in the data distribution included a morphological bias. This led to false negatives and bias against the detection of small grains. We used an expert model trained specifically on smaller pollen grains (a small grain detection model, SGDM) alongside GPDM detections to reduce taxonomic bias in our detection results (Fig. 3B). This is known as a mixture-of-experts approach (Nowlan and Hinton 1990).

Using the GPDM as our base model, we fine-tuned the SGDM detector by training exclusively on three small-grained taxa: Urticaceae-Moraceae (132 specimens for training/32 for validation), *Cecropia* (19/4), and Melastomataceae (25/4). We reserved examples from three other small-grained taxa – *Acalypha*, *Vallea*, and *Weinmannia* – as a held-out test set to assess the ability of the method to generalize to novel taxa with similar characteristics. The SGDM training set was a subset of the GPDM training set and the SGDM validation set was a subset of the GPDM validation set, ensuring a fair comparison for the two models, and trained the SGDM for 80 epochs.

We calibrated the SGDM confidence scores using a sigmoid function

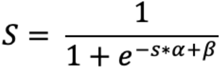

where *s* is the uncalibrated detection score, *α* and *β* are hyperparameters that we tuned experimentally, and *S* is the calibrated score (Platt 1999). The sigmoid function forces scores above the inflection point toward 1 and scores below the inflection point toward 0, and is a standard approach to adding nonlinearity to machine learning models. In our case, the function helps amplify small grain detections. We conducted several rounds of coarse-resolution testing within the range (0, 100) for *α* and (−10,10) for *β*, with step sizes of 1 or 10. After observing the general behavior of the curves, we varied *α* within the range (3.0, 6.0) with a step size of 0.1 and varied *β* within the range (3.0, 6.0) with a step size of 0.1. On the validation set, we found that the highest mean average precision (mAP) of the fused model was achieved at *α* = 5.5 and *β* = 4.9.

We fused detections from the GPDM and SGDM using NMS (IoU ≥ 0.3), eliminating overlapping detections and keeping only the detection with the highest score. We evaluated the performance of the fused model by comparing the precision-recall curve from the GPDM with that of the fused detector (GPDM + SGDM).

### Transfer Learning Across Imaging Domains

We next fine-tuned our GPDM, originally trained on PAL IV images, to new imaging domains, i.e., the PAL 1999 images. Because the taxonomic composition of the two cores were identical, differences between the two image datasets were primarily in the image resolution, color, and contrast. The PAL 1999 dataset included 300 image stacks from each of three slides; 80% were used as the training set and 20% as the validation set. The GPDM was trained for 120 epochs on PAL 1999 image stacks and annotations (Fig. 3C).

Although the PAL IV images had a resolution 0.146 mm per pixel and PAL IV images had a resolution of 0.225 mm per pixel, we did not rescale the images to simulate a more challenging domain gap. We evaluated the fine-tuned model performance by comparing the precision-recall curves for the slide scanner validation data before and after fine-tuning. To provide a performance baseline for comparison, we also trained a GPDM model trained from scratch on the PAL 1999 image stacks and annotations.

### Continual Learning with Human-in-the-Loop

We simulated a human-in-the-loop workflow that would allow us to train models when starting with incomplete data. Instead of training on all 900 images at once as in our transfer learning experiments, we split the images at the slide level into three sets of training and validation data to be used at three time periods (TP_0_, TP_1_, and TP_2_). We measured model performance as the decrease in the number of false positives and an increase in the recall rate. We manually verified true positives and false positives in our detection results from each time period and used these new labels to further fine-tune our models (Fig. 3D).

In TP_0_, we fine-tuned the GPDM on the TP_0_-training set and used the TP_0_-validation set to select the best-performing detector, which we called the TP_0_-detector. In TP_1_, we used the TP_0_-detector to help us annotate the TP_1_-training set. We ran the TP_0_-detector on the TP_1_-training set to get a set of detections. Next, we simulated human-in-the-loop verification of the detections by using the ground-truth annotations. If a ground-truth annotation and a predicted detection had an IoU >0.3, we considered the detection a true positive and included it in the set of cleaned annotations for the TP_1_-training set. Otherwise, the detection was considered a false positive and eliminated. We did not include missed detections in the cleaned annotations for the TP_1_-training set. We fine-tuned the TP_0_-detector on a training set composed of [TP_0_-training set + TP_1_-training set] and selected the best model using the TP_1_-validation set. We called this detector the TP_1_-detector. In TP_2_, we repeated the process of using the TP_1_ detector to annotate the current TP_2_-training set, then cleaning up the annotations. We fine-tuned the TP_1_-detector on a training set composed of [TP_0_-training set + TP_1_-training set +TP_2_-training set] and selected the best model using the TP_2_-validation set. We called this detector the TP_2_-detector.

### Zero-shot Segmentation

The shape of a pollen grain outline varies with orientation, preservation and species morphology. Not all pollen grains are circular in cross-section. As a result, circular detection masks can include extraneous background material, such as organic debris or adjacent grains. Equally, portions of the grain can be excluded when pollen shape deviates significantly from circular. New foundation segmentation models like Segment Anything Model-2 (SAM-2) allow segmentation of pollen grains without needing pollen images to finetune, i.e., “zero-shot segmentation” (Ravi et al. 2024). To evaluate the efficacy of SAM-2 with palynological images, we applied it to our pollen detections to produce more contoured segmentation masks that followed the true outlines of detected grains. We used the pollen grain center and the cropped pollen image output by our detection models as input for SAM-2. We also experimented with adding four points near the four corners of the detection mask as negative prompts to SAM-2 to unambiguously differentiate foreground from the background.

### Workflow Modules Development, Deployment, and Testing

We developed Command Line Interfaces (CLI) of the individual modules of the pollen analysis workflow to allow others to reproduce the analysis and adopt the image extraction and detection workflows. They are for tile cropping Hamamatsu NanoZoomer Digital Pathology Images (NDPI), running trained pollen image detection models, and post-processing with SAM-2 segmentation. Written in Python, the CLIs add additional functionalities like switching between running in serial and parallel mode, easy installation and dependency management using Docker or Apptainer on High Performance Computing (HPC) environments, access to optional parameters that can be passed on to the underlying software and to configure the runs, and logs to provide feedback to the user. We provide detailed developer-level documentation on installation and usage, and developed, deployed, and tested the CLIs using different environments (e.g., laptop, HPC, and Cloud).

## Results

### General Pollen Detection Model (GPDM)

We achieved a maximum detection recall rate of 93% and mean average precision (mAP) of 73.21% with the full PAL IV image dataset (Fig. 5A). A precision threshold of 20% removed the majority of false detections. At 20% precision, we achieved ∼90% recall across our experiments. The GPDM underdetected smaller, more transparent pollen grains such as Urticaceae-Moraceae, *Cecropia*, Melastomataceae, *Acalypha*, *Vallea,* and *Weinmannia* (Fig. 5C). For example, although *Cecropia* was a more common pollen type, with 19 training examples (Fig. 2M), none of the four images in the validation set were detected at the 20% precision level. In contrast, the larger and darker grains of *Polylepis* spp. (Fig. 2I), of which there were also 19 training examples, had 100% recall.

**Figure 5.**
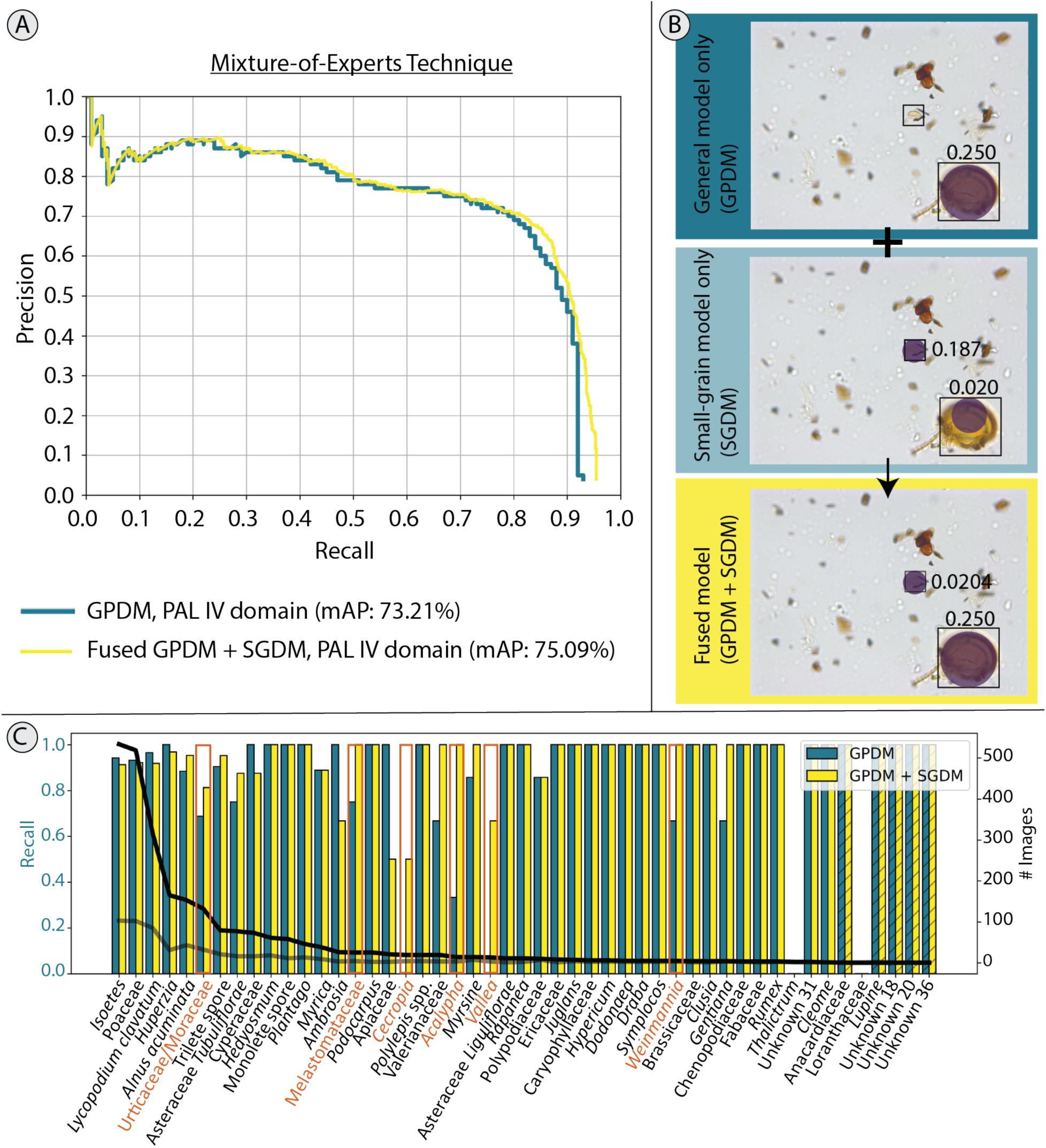
Results of GPDM, application of the mixture-of-experts technique, and application of transfer learning. (A) For the mixture-of-experts technique, comparison of the blue (GPDM) and yellow (fused model) curves shows how the addition of an expert model trained only on small pollen grains improves maximum model recall from 93% to 95%. (B) Comparison of detections from the general model, the small-grain model, and the fused model. Boxes in each panel indicate ground-truth labels and circles indicate detections. The color of the detections is arbitrary. Confidence scores are shown adjacent to each detection. SGDM confidence scores in the fused model have been calibrated as described in the text. A validation-set image stack containing a *Cecropia* pollen grain and *Isoetes* spore was fed into the two models. The GPDM detected the *Isoetes* spore with high confidence but missed the *Cecropia* grain. The SGDM was able to detect the *Cecropia* grain but had poor localization accuracy and low confidence for the *Isoetes* spore. The fused model kept both detections. (C) Blue and yellow bars indicate model recall by taxon for the GPDM and the fused (GPDM + SGDM) models respectively (left y-axis). The black and gray lines indicate the abundance distribution of taxa in the GPDM training and validation datasets, respectively (right y-axis). Both models were thresholded at 20% precision. The taxa highlighted in orange are particularly small-grained taxa, which have low rates of detection relative to the number of training examples. Hatched bars indicate taxa that had no training examples but were in the validation set and detected by the detector.

### Mixture-of-Experts Technique

Fusing the SGDM with the GPDM increased maximum recall by 2% and increased mAP from 73.21% to 75.09% (Fig. 5A). Recall at the 20% precision level increased for Urticaceae-Moraceae from 69% to 81%, Melastomataceae from 75% to 100%, and *Cecropia* from 0% to 50% (Fig. 5C). *Acalypha*, *Vallea*, and *Weinmannia*, the three remaining taxa with small, transparent pollen grains, also showed improvement, despite not being included as SGDM training examples (Fig. 5C). The fused SGDM + GPDM model shows an increase in model precision within a large range of recall in (0.1, 0.9), indicating that the SGDM added some high-confidence small-pollen detections (Fig. 5A). The increase in maximum recall shows that the SGDM added detections that are missed by the GPDM (Fig. 5A, 5B).

### Transfer Learning Across Imaging Domains

We achieved an mAP of 32.56%, maximum recall of 82%, and 10% precision at the 80% recall level when we forward-passed the PAL 1999 (slide scanner) images through the GPDM trained on PAL IV (upright microscope) images. The low performance demonstrated the extent to which the detection models were sensitive to domain differences caused by illumination, optical resolution, and camera property offsets. After fine-tuning the model with 900 PAL 1999 training images, we achieved an mAP of 65.99% and precision increased from 10% to 46% at the 80% recall level (Fig. 6). Maximum recall increased from 82% to 93%. The model trained from scratch on the 800 PAL 1999 images achieved an mAP of only 59.27% (Fig. 6).

**Figure 6.**
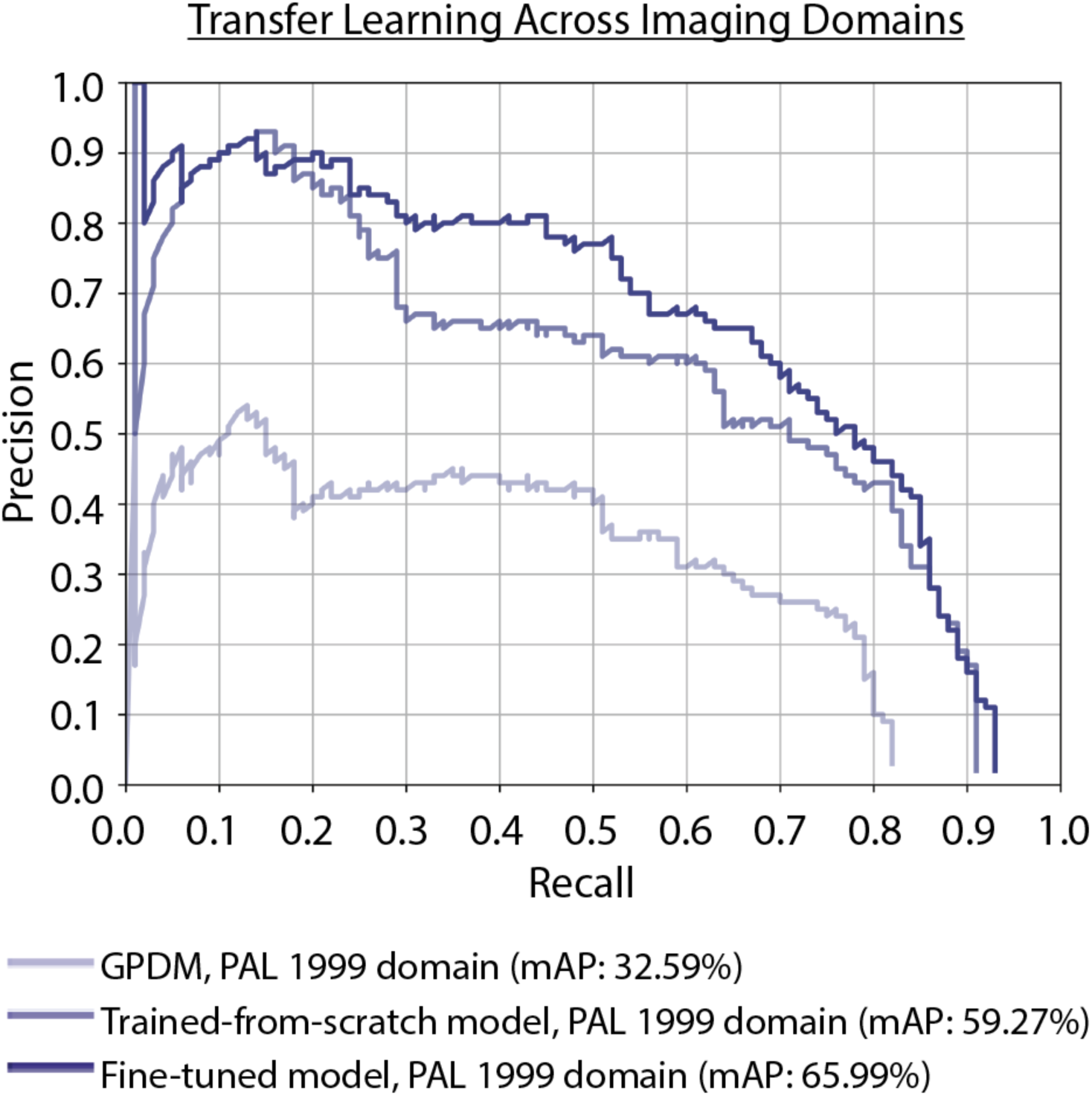
Comparison of the lightest and darkest purple curves shows the improvement of the model after fine-tuning. Maximum recall increased from 82% to 93% and precision increased from 10% to 46% at the 80% recall level. The medium purple curve, representing the performance of a model trained from scratch on the PAL 1999 domain, shows that training from scratch on a small dataset is not as effective as fine-tuning.

### Continual Learning with Human-in-the-Loop

Human-in-the-loop fine-tuning increased mAP with each iteration, from 34.11% to 58.72% in TP_0_, from 48.25% to 62.60% in TP_1_, and from 61.57% to 76.01% in TP_2_ (Fig. 7). To compare model performance across all three time periods, we also evaluated each model on a common validation set (composed of the TP_0_+TP_1_+TP_2_ validation sets). Against this common validation set, mean average precision increased from 41.10% with the TP_0_ model to 59.93% using the TP_2_ model (Fig. 7D).

**Figure 7.**
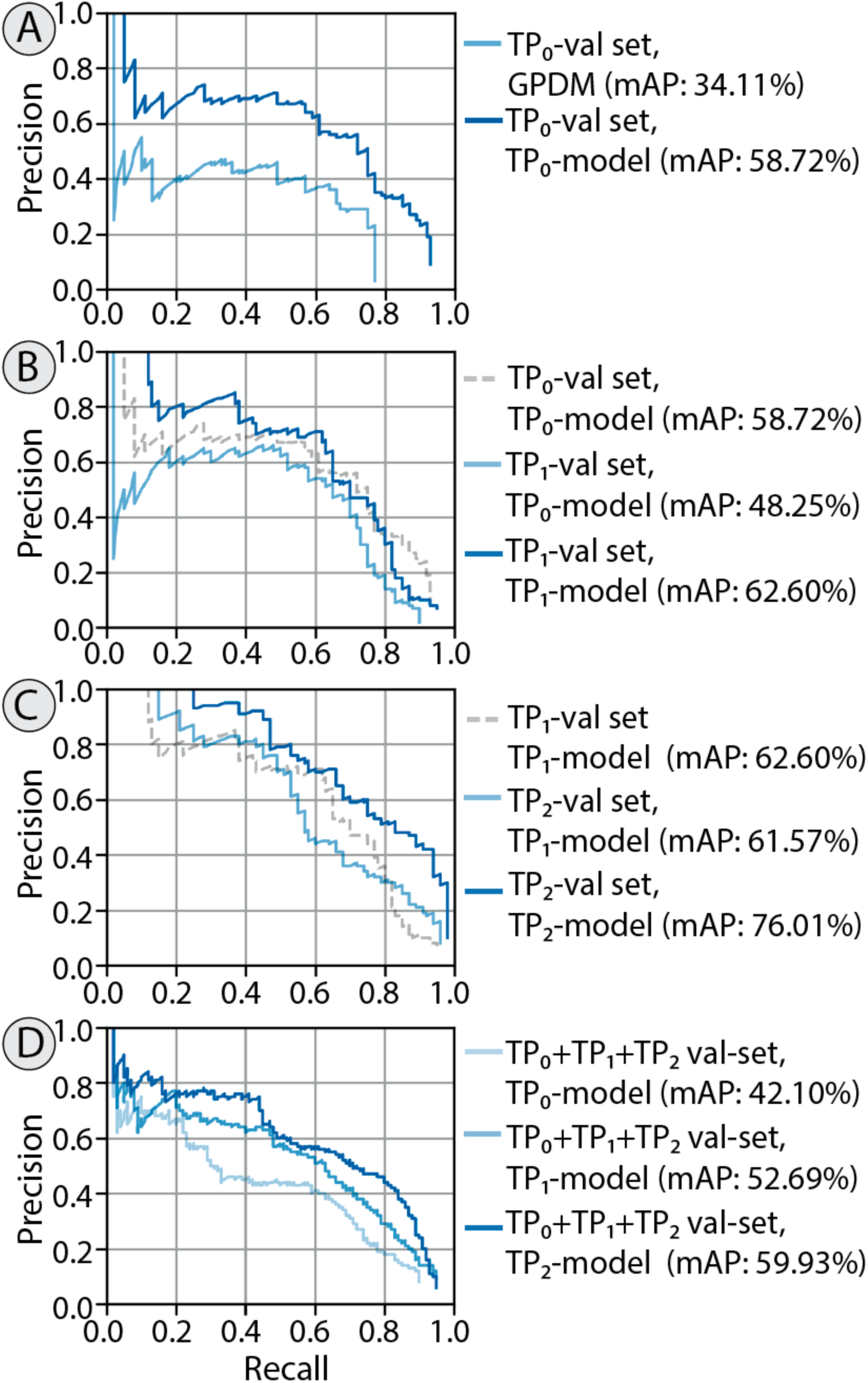
Panels (A - C) show the improvement in model performance from fine-tuning in three time periods. (A) The light blue curve represents the performance of the GPDM on the TP_0_ validation set. The darker blue line shows the increase in model performance after fine-tuning on the TP_0_ training set with ground-truth annotations. (B) depicts the second time step, TP_1_. The dashed gray curve is the same as the dark blue curve in (A). Comparing the dashed gray curve to the light blue curve shows the drop in model performance when switching slides and introducing a domain gap. Comparing the light and dark blue curves, we see that fine-tuning on the expert-verified TP_0_ model detections of the TP_1_ training set improves model performance again. (C) A similar pattern is shown in the next time step, TP_2_. (D) A comparison of the performance of the three models on the same validation set. Here, the TP_0_+TP_1_ +TP_2_ validation set.

### Zero-shot Segmentation

The SAM-2 segmentation masks with the highest probability scores closely followed pollen grain shape (Fig. 8). We used a single point, the center of the original detection bounding box, to prompt SAM-2. Because the detection bounding box tightly bounds the detection, the center of the detection mask identified the region of interest and provided a clean input prompt for SAM-2. Adding four points near the four corners of the detection mask as negative prompts did not improve the results. This may be because the image background significantly differs from the pollen. However, in situations where the foreground and background have less color difference, this approach could be used. Notably, SAM-2 was able to recognize nearly the entirety of a pollen grain although the foundation model had not been trained on pollen images. To completely capture the pollen wall, we applied dilation (an image processing morphological operation; 5 x 5 kernel over 1 iteration) to expand the area captured by the segmentation mask and ensure that the mask completely captured the pollen wall and external ornamentation.

**Figure 8.**
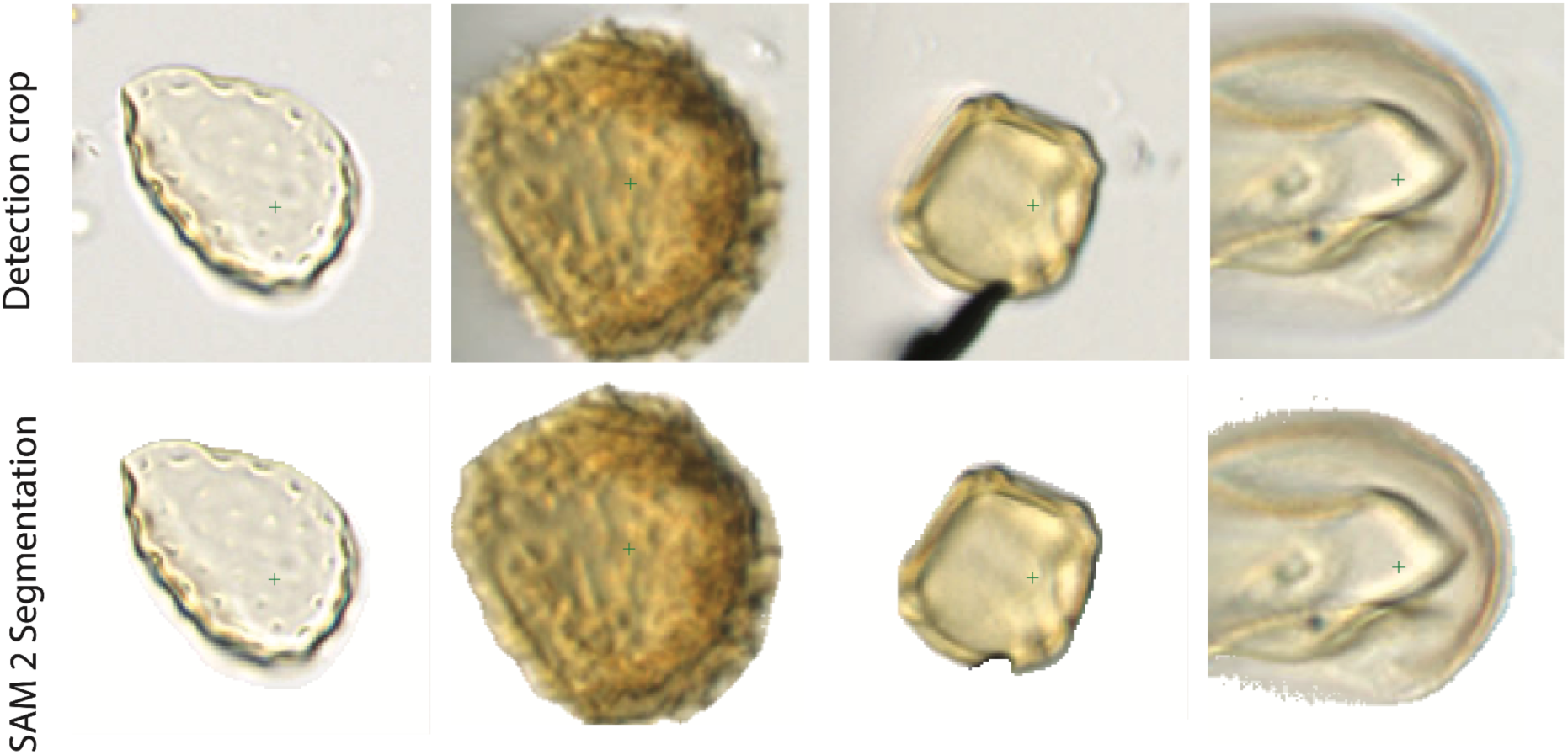
Illustration of segmentation of cropped and detected pollen images using SAM-2. Top row shows the original cropped detection. Bottom row shows the SAM-2 segmentation results.

## Discussion

Rapid slide imaging is possible with the commercial availability of automated stages and slide scanning microscopes (e.g., Punyasena et al. 2022; Theurkauf et al. 2023). Efficient object detection in scanned slide images will streamline the process of annotating and curating training data and effective, generalized detectors will lower barriers to the adoption of machine learning in palynology and other fields of microfossil research.

Several studies have developed detection and classification methods for analyzing environmental pollen on slides (e.g., Gimenez et al. 2024; Li et al. 2024; Tešendić et al. 2022; Battiato et al. 2020; Kubera et al. 2022; Punyasena et al. 2022; for a review see Buters et al. 2022), and there are commercially available instruments for detecting and classifying airborne pollen (Oteros et al. 2020). Additionally, imaging flow cytometry can process particles rapidly and some instruments are capable of simultaneously taking brightfield and fluorescence images of particles. The autofluorescence of pollen grains are used to visually isolate them (Dunker et al. 2021; Barnes et al. 2024).

However, fossil pollen detection presents its own unique challenges. Pollen morphology is altered by the processes of fossilization, introducing taphonomic artifacts and morphological distortions that vary with taxonomic composition and depositional environments (Cushing 1967; Delcourt and Delcourt 1980; Campbell 1999). Fossil samples may also contain more inorganic sediment, a larger number of unknown plant taxa, and a greater diversity of non-pollen organic debris, all of which introduce noise to automated analyses. Most fossil pollen material is mounted on slides. Automated detection methods compatible with traditional counting and archival material rely on scanning slides using multiple focal planes (12 to 5 planes, spaced <3 to 8 um; Von Allmen et al. 2024; Theuerkauf et al. 2024; Gimenez et al. 2024).

While previous studies fused the sharpest portions of the image stacks into a single 2D image using focus stacking algorithms, the detection model described in our analysis is trained directly on an unfused image stack. This approach retains the spatial information contained in all focal planes and also allows post-hoc selection of the most in-focus plane or the entire image stack for downstream analyses. We use the mAP metric to evaluate and benchmark model performance because it is agnostic to IoU and confidence thresholds. That is, at different IoU and confidence thresholds, recall and precision will differ, but by using precision-recall curves we can evaluate the relationship between the two metrics at all confidence thresholds. The confidence threshold can be set by the user depending on the application. In most cases, a high recall rate is preferable because false detections can be further eliminated during classification or manual post-processing (von Allmen et al. 2024; Theuerkauf et al. 2024).

The choice of CNN architecture can vary. We used ResNet34 (He et al. 2016). Von Allmen et al. (2024) used CenterNet Hourglass104 (Padilla et al. 2019). Theuerkauf et al. (2023) used Faster R-CNN (Ren et al. 2016). Gimenez et al. (2023) used the joint detection and classification model YOLOv5 (Zhang et al. 2022). These previous studies achieved detection accuracies >90%, but with a limited number of taxa (10 - 11 pollen types). They demonstrate that while high recall rates are possible within a small number of taxa, the diversity and long-tail distribution of pollen taxa within paleontological samples makes detecting rare and unusual morphotypes difficult. Theuerkauf et al. (2023) addressed this imbalance by under-sampling majority classes and von Allmen et al. (2024) by selectively balancing training data.

We also found that the GPDM under-detected taxa with small, transparent grains: Urticaceae-Moraceae, *Cecropia*, Melastomataceae, *Acalypha, Vallea,* and *Weinmannia* (Fig. 5C). These smaller-grained taxa were not only rare, but also differed morphologically from the majority of the training data. In contrast, rare taxa that had a similar size, shape, and color to taxa in the training set were easily detected by the GPDM. For example Ericaceae, *Rumex*, and *Lupine* were detected with 100% recall in the test set (Fig. 5C), despite having ten or fewer training examples.

The mixture-of-experts technique (Nowlan and Hinton 1990) allowed us to address taxonomic bias without additional annotated data. Fusing an expert small-grain detection model (SGDM) with the GPDM increased detections of rare, small-grained taxa. Three taxa that were not included as SGDM training examples (*Acalypha*, *Vallea*, and *Weinmannia*) also showed improvement (Fig. 5C). This suggests that the SGDM-learned features were transferable across taxa with similar morphologies. However, taxa with few training examples can still be missed stochastically based on which images were included in training and validation. For example, two rarer taxa, Loranthaceae and *Thalictrum*, were entirely missed by both the GPDM and SGDM (Fig. 5C). At 20% precision, we achieved ∼90% recall across our experiments. Lowering the precision threshold increased detections and recall, but introduced a greater percentage of false detections. Threshold selection, therefore, is ultimately dependent on the application and tolerance for missed versus false detections and will likely need to be determined on a case-by-case basis. Previous work has demonstrated that the abundance of a particular pollen taxon, the abundance of a particular pollen morphology, and the degree of morphological similarity to organic debris all contribute to detection errors (Gimenez et al. 2024; Theuerkauf et al. 2023). Including reference specimens in model training is a potential solution, but as Durand et al. (2024) show, models trained on fresh reference specimens may not be well-suited for fossil pollen analysis.

Differences in sample preparation and imaging practices create domain gaps among image datasets. Previous work has shown that transfer learning is effective in pollen classification (Rostami et al. 2023; von Allmen et al. 2024), and we demonstrate that transfer learning is also effective in the context of detection. This allows us to deal with domain gaps introduced by different microscope models and scanning conditions. Detection models need to work with multiple imaging sources as standardizing imaging across large imaging projects is not always feasible, manual annotations are costly in terms of expert time, and training new models for every dataset is not always possible. However, it is possible to fine-tune pre-trained detection models on relatively small amounts of training data from new imaging domains. We needed to annotate only 5% of images from a new imaging and sampling domain to increase mAP from 32.56% to 65.99% (Fig. 6), while training from scratch on the new domain alone achieved an mAP of 59.27% (Fig. 6). Collaboration between institutions with different microscopes and imaging equipment is therefore a feasible (and potentially desirable) approach, as diverse training data will aid in the development of large, robust pollen analysis pipelines.

With slide scanners that can scan an entire slide in under one hour and detection and classification models that can be trained in days (e.g., Punyasena et al., 2022; Theuerkauf et al., 2023), annotation remains the primary bottleneck in automated pollen analysis workflows. Continual learning – with human-in-the-loop annotations and iterative fine-tuning – allows models to improve as new images and new data are introduced. Confidence scores can be used to efficiently annotate false detections manually or using semi-automated methods. New annotations can be used to further fine-tune models, and these models can improve over time with expert feedback on model results. Strategies for effective human-in-the-loop data annotation are worth studying as a stand-alone problem, as for instance Gimenez et al. (2024) has done for studying the effect of labeling specificity on model performance.

Finally, as foundation segmentation models like SAM-2 become incorporated into analysis workflows, procedures for detection and segmentation will become further streamlined. These foundation models can efficiently segment objects within an image with a single prompt (e.g., a mouse click, a coarse bounding box, or a coarse mask obtained from an upstream task, etc.), producing clean segmented images for training classification models. However, for palynological samples, trained detection models may still be needed to distinguish pollen and other specimens of interest from the diverse organic material that can be found on a palynological slide. However, once detected, foundation segmentation models are able to use a single set of coordinates to produce the clean contoured images needed for machine learning classification analyses.

## Conclusions

Working in open-world scenarios means that we do not know *a priori* the full diversity that we will encounter in our analyses, so we need methodological approaches in which we continually improve both detectors and classifiers. Mixture-of-experts techniques, domain fine-tuning, and continual learning allow researchers to build upon shared generalized machine learning models, allowing each new generation of machine learning models to become more powerful than the last. Mixture-of-experts technique allows us to build models that are more resilient to changes in data distribution. Fine-tuning allows us to efficiently apply general models to new domains. These general models will also help annotate new data in new domains, using human-in-the-loop annotation to efficiently build large and diverse pollen image datasets. Efficient object detection in scanned slide images will streamline the process of annotating and curating training data and effective, generalized detectors will lower barriers to the adoption of machine learning in palynology and other fields of microfossil research. While the underlying architecture of machine learning models may change, incorporating open-world methods will result in flexible and adaptable workflows that are accessible to all paleobiologists.

## Authors’ Contributions

JTF and SWP led the writing of the manuscript, with contributions from all authors. SWP, SK, JTF, and THD conceived the ideas and designed the methodology. JTF, SPS, and SK wrote the code. SPS deployed and performed initial testing of the CLIs in an HPC environment (Delta) and a Cloud environment (Radiant) all hosted at the National Center for Supercomputing Applications (NCSA). THD led the collection and preparation of the palynological samples and verified the pollen identifications. JTF prepared the PAL1999 samples, imaged the PAL1999 slides, and provided expert palynological annotations for the PAL1999 image stacks. SWP and SK directed the research.

## Acknowledgements

Thanks to Kimberley Hagemans (formerly at Utrecht University) and Thya van den Berg (University of Hull, formerly at Utrecht University) for preparing, annotating, and imaging the PAL IV samples; Giovanni Dammers (Utrecht University) for his supervision in preparing the samples; David Tcheng (formerly at the National Center for Supercomputing Applications, University of Illinois) for developing the Matlab annotation script used for the PAL IV samples; and Edwin Bennink (Utrecht University) for modifications and updates to the annotation script. Funding for this research was provided by the University of Illinois Campus Research Board (Grant RB22079 to SWP), the University of Illinois School of Integrative Biology Francis M. and Harlie M. Clark Research Support Grant to JTF, the Dutch Research Council (NWO) (Grant 824.14.018 to THD), and the Institute of Collaborative Innovation and the University of Macau (Grant SRG2023-00044-FST to SK). This research was also supported in part by the Illinois Computes project through the University of Illinois Urbana-Champaign and the University of Illinois System. This work used the National Center for Supercomputing Applications’ (NCSA) Delta computing platform, through allocation EES240072 from the Advanced Cyberinfrastructure Coordination Ecosystem: Services & Support (ACCESS) program, which is supported by U.S. National Science Foundation grants #2138259, #2138286, #2138307, #2137603, and #2138296 and the NCSA private cloud computing service Radiant. We thank Associate Editor Caroline Stromberg and three anonymous reviewers for feedback that substantially improved the manuscript.

## Data availability

Image stacks, annotation data, and trained detection models are archived through the Illinois Data Bank (Feng et al. 2025). We developed Python-based CLIs that are containerized using Docker for easy installation and provide many features like serial and parallel processing modes, storing output in different forms (folders or zip files), optional parameters for customizing the runs of the applications and for interacting with the underlying software. The developed CLIs were successfully tested on MacOS (Apple M Chips and Apple Intel Chips) and NCSA’s Delta High Performance Computing (HPC) system using Apptainer. Source code for the described analysis is available on GitHub under Apache 2.0 open-source license at https://github.com/paleopollen/ndpi-tile-cropper-cli, https://github.com/paleopollen/pollen-detection-cli, and https://github.com/paleopollen/segment-anything-2. The original set of Jupyter Notebooks are available at(github.com/paleopollen/open_world_pollen_detection).

